# The “creatures” of the human cortical somatosensory system

**DOI:** 10.1101/675611

**Authors:** N. Saadon-Grosman, Y. Loewenstein, S. Arzy

## Abstract

Penfield’s description of the “homunculus”, a “grotesque creature” with large lips and hands and small trunk and legs depicting the representation of body-parts within the primary somatosensory cortex (S1), is one of the most prominent contributions to the neurosciences. Since then, numerous studies have identified additional body-parts representations outside of S1. Nevertheless, it has been implicitly assumed that S1’s homunculus is representative of the entire somatosensory cortex. Therefore, the distribution of body-parts representations in other brain regions, the property that gave Penfield’s homunculus its famous “grotesque” appearance, has been overlooked. We used whole-body somatosensory stimulation, functional MRI and a new cortical parcellation to quantify the organization of the cortical somatosensory representation. Our analysis showed first, an extensive somatosensory response over the cortex; and second, that the proportional representation of body-parts differs substantially between major neuroanatomical regions and from S1, with, for instance, much larger trunk representation at higher brain regions, potentially in relation to the regions’ functional specialization. These results extend Penfield’s initial findings to the higher level of somatosensory processing and suggest a major role for somatosensation in human cognition.

## Introduction

The establishment of the homunculus, a schematic drawing reflecting the disproportional representation of the parts of the human body on the motor and somatosensory cortex, was an important milestone for the neurosciences. Eighty years ago Penfield and Boldrey electrically stimulated the cortical surface of patients undergoing brain surgery. They used the patients’ subjective reports of somatosensory sensations to identify representation of different body-parts in the postcentral gyrus, known as the primary somatosensory cortex (Penfield and Boldrey, 1937). One of their significant findings was that the cortical surface area associated with a body-part (“spatial distribution” of body-parts) is not proportional to the surface area of the body-part itself. Instead, the face and hand are overrepresented, while the trunk and leg occupy disproportionately small cortical areas. A schematic drawing of a human figure with body-parts corresponding to the size of their cortical (S1) representation, yielded the famous “grotesque creature” with large hand and lips and rather small trunk and leg, known as the somatosensory homunculus (Penfield and Boldrey, 1937; Penfield and Rasmussen, 1950; Penfield and Jasper, 1954). Later, it was suggested that the spatial distribution depicted by the homunculus reflects the level of neuronal peripheral innervation (Woolsey et al., 1942; Sur et al., 1980; Catani, 2017). The disproportionate spatial distribution of body-parts is likely to be of functional significance, underlying the greater somatosensory discrimination ability for the enlarged body-parts in the corresponding cortical homunculus.

Penfield and his colleagues also identified somatosensory representations in other cortical areas. These include the precentral gyrus (the primary motor area, M1; (Penfield and Boldrey, 1937)), the superior bank of the lateral fissure (the secondary somatosensory area, S2), the insular cortex, and the medial cortex (supplementary motor area, SMA; (Penfield, 1950; Penfield and Jasper, 1954; Penfield and Faulk, 1955)). Notably, these representations were less grounded. Later studies used electrophysiological recordings in non-human primates and functional neuroimaging in humans in response to somatosensory stimulation to further identify and characterize the cortical somatosensory system (e.g., Kaas et al., 1979; Fox et al., 1987; Burton et al., 1993; Lim et al., 1994; Nakamura et al., 1998; Kaas and Collins, 2001; Ruben et al., 2001; Fitzgerald et al., 2004; Mazzola et al., 2005). Additional somatosensory representations were detected in the superior and inferior parietal lobules (Sakata et al., 1973; Ruben et al., 2001; Young et al., 2004; Huang et al., 2012), inferior frontal gyrus, frontal operculum (Hagen et al., 2002) and the cingulate cortex (Arienzo et al., 2006). Large body of works focused on specific localizations of body-parts representations across the different cortical regions. In contrast, surprisingly little attention has been given to the spatial distribution of body-parts across the cortical surface, the property that gave Penfield’s homunculus of S1 its famous characteristic “grotesque” appearance.

The goal of this research is to quantify the spatial distribution of body-parts across the entire somatosensory cortex. We first characterized the somatosensory cortex by measuring its response to a bilateral whole-body continuous tactile stimulation, using functional magnetic resonance imaging (fMRI). Interestingly, our measurements revealed extended somatosensory cortical response to tactile stimulation. Second, we quantified the spatial distribution of body-parts in different anatomically-distinct cortical regions. Spatial distribution of body-parts substantially varied in between regions and S1. We interpret the differences in these distributions as reflecting functional specialization.

## Materials and Methods

### Participants

20 healthy participants (9 females, age: 27.45±3.33 year-old (mean±SD)) participated in the study. Participants did not report any history of neurological, psychiatric, or systemic disorder. All participants gave written informed consent, and the study was approved by the ethical committee of the Hadassah Medical Center.

### Experimental Paradigm

Whole-body continuous periodic brush movement was applied from lips-to-toes and from toes-to-lips in two different scanning runs (Fig. S1A; (Saadon-Grosman et al., 2015; Tal et al., 2016)). Each scanning run included 14 whole-body stimulation cycles, equally divided between the two body-sides (counter-balanced between participants). The length of each stimulation cycle was 15 s, which was followed by a 12 s of rest (baseline). Experimental runs started with 30 s of rest before the first cycle onset and ended with 4.5 s after the last cycle offset, in addition to 12 s of rest between body sides. Natural light-touch stimulation was delivered using a 4-cm-wide paint-brush (with extended handle of 0.65 meter plastic stick) by the same experimenter, who was well-trained before the scans to maintain a constant pace and pressure during the runs. The experimenter wore fMRI-compatible headphones, delivering preprogrammed (Presentation; Neurobehavioral Systems) auditory cues signaling the precise timing of the body-part sequence, which enabled a controlled velocity of tactile stimuli.

### Functional MRI Image Acquisition Procedures and Preprocessing

All participants were scanned at the same site using a Siemens Skyra 3T system (32-channel head coil) with the same imaging sequence. Blood oxygen level dependent (BOLD) fMRI was acquired using a whole-brain, gradient-echo (GE) echoplanar (EPI) [repetition time (TR)/time echo (TE) = 1,500/27 ms, flip angle = 90, field of view (FOV) = 192 × 192 mm, matrix = 64 × 64, 26 axial slices, slice thickness/gap = 4 mm/0.8 mm. In addition, high resolution (1 × 1 × 1 mm) T1-weighted anatomical images were acquired to aid spatial normalization to standard atlas space. The anatomic reference volume was acquired along the same orientation as the functional images [TR/TE = 2,300/2.98 ms, matrix = 256 × 256, 160 axial slices, 1-mm slice thickness, inversion time (TI) = 900 ms]. Preprocessing was performed using the Brain Voyager QX 20.4.0.3188 software package (Brain Innovation) and NeuroElf (http://neuroelf.net), including head motion correction (trilinear interpolation for detection and sinc for correction), slice scan time correction, and high-pass filtering (cutoff frequency, two cycles per scan). Temporal smoothing (FWHM = 4 s) and spatial smoothing (FWHM = 4 mm) were additionally applied (Saadon-Grosman et al., 2015). Functional and anatomical datasets for each participant were co-registered and normalized to standardized MNI (ICBM-152) space. All further analyses were performed using in-house custom Matlab (Mathworks, Inc.) scripts.

### Cross-Correlation Analysis

To identify the cortical distribution of the somatosensory system we used a cross correlation analysis. A boxcar function (3 s) was convolved with a two gamma hemodynamic response function (HRF), to derive a predictor for the analysis. This predictor and the time course of each voxel were cross-correlated to measure responses to different parts of the stimulation cycle (body-parts). The first volume in the stimulation block was excluded to avoid any effects of expectancy or surprise. The predictor was cross correlated across all volumes in the block except the last to allow averaging of the two opposite stimulation directions (“start lips” and “start toes”). Stimulation duration of each cycle had 10 volumes, thus, cross correlation analysis produced 8 correlation values for each voxel, indicating correlation to different parts of the stimulation cycle (each volume was assigned to a specific body part by its stimulation time: 1-lips, 2-distal upper limb, 3-proximal upper limb, 4-upper trunk, 5-lower trunk, 6-proximal lower limb, 7-mid lower limb, 8-distal lower limb). For each voxel of each participant, we flipped the order of the correlation values of the start toes paradigm and then averaged the correlation distributions of both start lips and start toes directions. The averaged distribution maximum defined the preferred body-part (lag value) of each voxel. A correlation threshold of r > 0.251 was applied to identify voxels responding significantly to the stimulation (t-test, α = 0.05, Bonferroni corrected for multiple correlations; 8 lag values × two paradigms, 134 degrees of freedom, p<0.003). To generate a group map, all correlation distributions across participants were averaged in each voxel. Only voxels that were above the significance threshold in more than 2/3 of the participants were included in the analysis (random effect yielded similar results, Fig. S2).

### Cortical parcellation and identification of somatosensory responsive areas

In this study we applied a recently introduced multi-modal data-driven parcellation (Glasser et al., 2016). This parcellation uses multimodal magnetic resonance images from the Human Connectome Project (HCP) and an objective semi-automated neuroanatomical approach to delineate 180 areas per hemisphere, bounded by sharp changes in cortical architecture, function, connectivity, and/or topography (Glasser et al., 2016). Since this parcellation is surface based, cross-correlation maps and voxel time courses were projected on an averaged inflated cortical surface (Trilinear interpolation, data in depth along vertex normal from -1mm to 3mm of grey-white matter border; FreeSurfer’s, fsaverage template brain; (Desikan et al., 2006)). Parcellation areas containing more than 50% of vertices responding to body stimulation were defined as somatosensory responsive areas (Table S1). Only significant vertices were considered, this threshold was used to eliminate areas which did not pass majority rule (i.e. areas with less than 50% significant vertices). Yet, we did not include insignificant vertices in the analysis, even in supra-threshold parcellation areas.

### Gross-anatomy classification

The somatosensory system as found here lies over four gross anatomical regions, including the anterior part of the parietal lobe (from BA 3a to the ventral and medial intraparietal areas (BA 7)), the posterior part of the frontal lobe (from BA 4 to the anterior end of BA 6), the superior part of the medial wall (from the medial end of the pre and post central gyri to the middle cingulate gyrus) and the operculum-insular cortex (from parietal and frontal operculum to temporal operculum through the posterior insula). In order to define these regions precisely we utilized the above mentioned parcellation. We classified each of the somatosensory parcellation areas according to the following criteria: the parietal region includes all areas on the lateral surface posterior to the central sulcus, the frontal region includes all areas on the lateral surface anterior to the central sulcus, the medial region includes all areas on the medial wall, and the operculum-insula region includes areas inferior to the parietal and frontal lobes of the operculum and insular cortex (Table S1, “Neuroanatomical results for a multimodal parcellation of human cerebral cortex” (Glasser et al., 2016)). Note that areas 4, 6mp and 7Am (Glasser et al., 2016) on the lateral surface extend into the medial wall and that area 5L as noted in the parcellation is somehow different from Brodmann’s original definition thus is part of the medial region. Additional parcellation areas that were not included in these anatomical regions as part of the spatial continuous representation were not considered in the analysis (eight in the right hemisphere and seven in the left hemisphere, see Table S1). Relative proportions of each body-part within the entire somatosensory system and in specific anatomical regions (gross and parcellation areas) were calculated according to the percentage of significant vertices with a given lag value (lips-1, upper limb-2,3, trunk-4,5, lower limb-6,7,8). To estimate the level of confidence we applied bootstrapping by resampling participants to create 1,000 cross correlation maps, errors are represented by the standard deviation of body-parts proportions. In addition, we created a schematic drawing of the relative proportions of body parts by modifying Penfield’s S1 homunculus drawn by “Cortical Homunculus”, The Homunculus mapper, license CC BY 2.0 (https://www.maxplanckflorida.org/fitzpatricklab/homunculus/). Body parts of S1 schematic drawing were rescaled *(using Adobe Illustrator*^*®*^ *CS6)* following the results presented in Fig. S4. The body parts were then reconnected into a modified homunculus schematic drawing. Note that not all body-parts were stimulated (e.g. tongue and forehead). Therefore, these body-parts are represented according to the schematic figure of S1 homunculus, similar to the illustration by Penfield and colleagues (Penfield and Boldrey, 1937).

### Statistical testing of differences in body-parts spatial distribution

When comparing the spatial distribution of body parts between two regions, we spatially permuted the relevant vertices 1000 times and recomputed the distribution of body parts in the two regions. Significance was tested by computing the fraction of permutations, in which the difference between the proportions of the corresponding body parts in the two regions was larger than the experimentally-measured.

## Results

### Spatial distributions of body-parts across the somatosensory responsive cortex

For each participant, we identified cortical voxels that responded significantly to the somatosensory stimulation (p<0.05, Fig. S1). Overall, group map shows that somatosensory representation in the contralateral hemisphere (which we operationally define as the somatosensory responsive cortex; ipsilateral maps, Fig. S3) spans more than 30% of the cortical surface (35.7±5.1% of the right hemisphere and 30.4±5.8% of the left hemisphere, surface area). The somatosensory map includes the previously identified somatosensory regions in S1, M1, S2, SMA, posterior insular cortex, superior and inferior parietal lobules, inferior frontal gyrus, frontal operculum and the cingulate cortex (Fig. 1A).

**Fig. 1.**
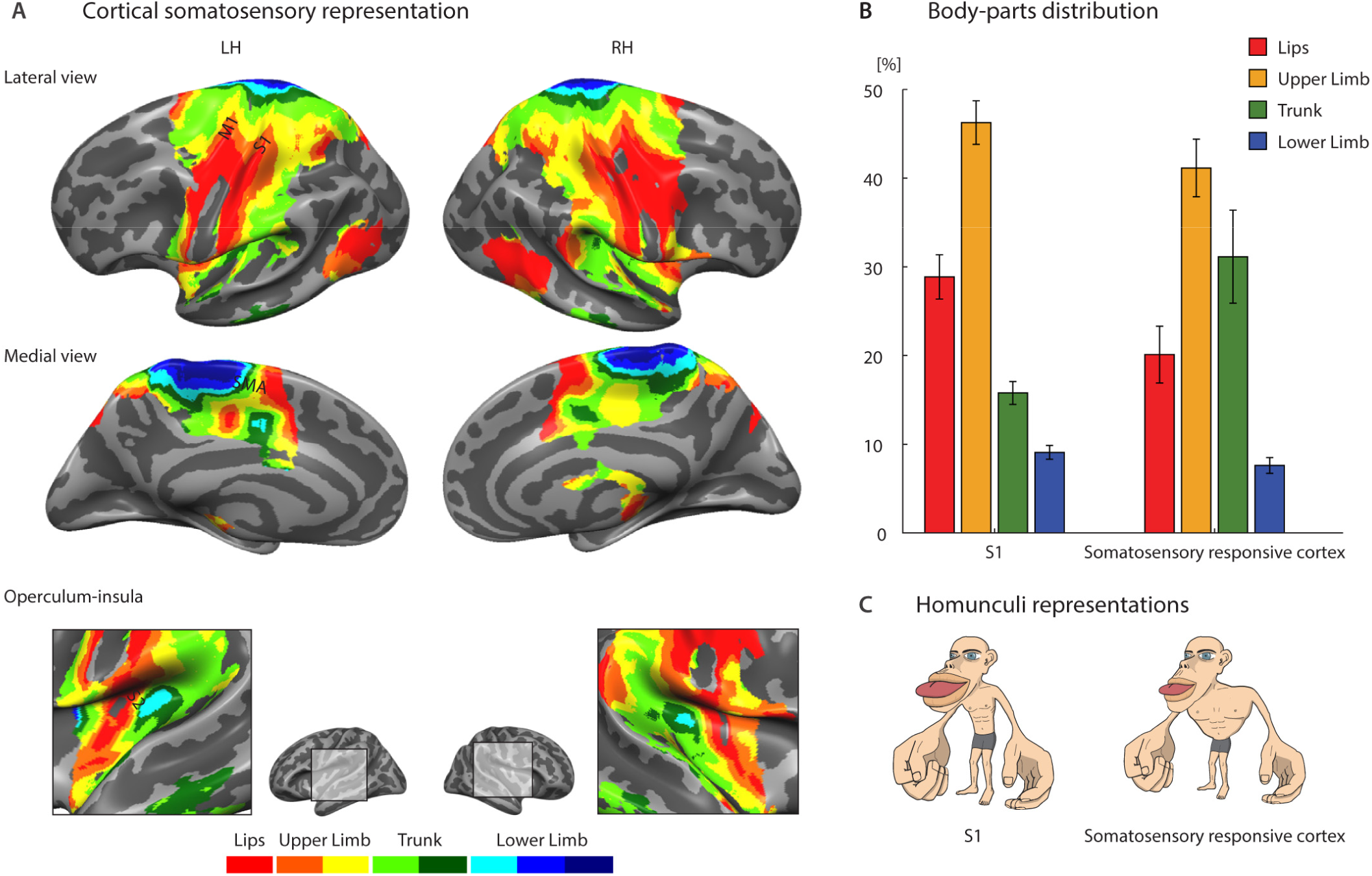
Somatosensory cortical representation and body-parts spatial distribution. **(A)** Cross correlation group maps (N=20; single participants) corresponding to stimulation of the contralateral body side are shown on the lateral and medial surfaces as well as the operculum and insula cortices (LH: left hemisphere; RH: right hemisphere; level of significance: at least 2/3 of the participants with significance p<0.05; random effect yielded similar results, Fig. S2). Color code represents body-parts: lips (red), distal upper limb (orange), proximal upper limb (yellow), upper trunk (light green), lower trunk (green), proximal lower limb (light blue), mid lower limb (blue) and distal lower limb (dark blue). Landmarks: M1: primary motor area; S1: primary somatosensory area; S2: secondary somatosensory area; SMA: supplementary motor area. **(B)** Quantification of body-parts [lips, upper limb (proximal and distal combined), trunk (upper and lower combined) and lower limb (proximal, mid and distal combined)] spatial distribution in S1 (left, BA: 3a,3b,1,2) and in the entire somatosensory responsive cortex (right) averaged across two hemispheres (Error bars are standard deviation, computed by bootstrapping over participants). Note the different percentage of trunk and lips between S1 and the entire somatosensory responsive cortex. **(C)** The original schematic homunculus representation of S1 adopting Penfield’s homunculus (left) and a modified version for the entire somatosensory responsive cortex (right).

The quantification of the spatial distribution of body-parts in S1 (defined according to Brodmann’s areas (BAs) 1, 2, 3a, 3b (Glasser et al., 2016)) was found to be comparable to Penfield’s somatosensory homunculus (Penfield and Boldrey, 1937; Catani, 2017). That is, extensive representation of the upper limb (46.3±2.5%) and lips (28.9±2.6%), and lesser representation of the trunk (15.8±1.3%) and lower limb (9.1±0.8%; Fig. 1B, left). However, when we quantified the entire somatosensory cortex, the spatial distribution was significantly different. While the fractions of the cortical area devoted to the upper (41.1±3.3%) and lower (7.6±0.9%) limb are both slightly (but significantly) smaller than those in S1 (p<0.001), the area devoted to the lips (20.1±3.3%) is only 70% of that in S1, and the trunk area (31.1±5.3%) is almost double than in S1 (both significantly different, p<0.001). Adopting Penfield’s homunculus representation, S1 and the entire somatosensory responsive cortex are represented by very different homunculi (Fig. 1C). These differences imply that principles other than the level of neuronal peripheral innervation, which is believed to underlie this distribution in S1 (Woolsey et al., 1942; Sur et al., 1980; Catani, 2017), determine the spatial distribution of body-parts in the cortex as a whole.

To study these principles, we asked whether anatomically-distinct regions are characterized by different spatial distributions. According to fundamental subdivisions of the cortex’ gross anatomy, somatosensory representations are found within the parietal lobe (S1 and posteriorly), the frontal lobe (anterior to S1), the medial wall (medial to S1) and operculum and insula (inferior to S1) regions. To precisely segment the cortex into these four regions, we capitalized on a recently-introduced, data-driven multi-modal functional cortical parcellation that divides each hemisphere into 180 anatomically and functionally defined areas (Glasser et al., 2016). By using these areas to draw the borders between the four regions (Fig. 2A), we were able to associate each vertex with its corresponding region (see also Table S1 and Fig. S5A).

**Fig. 2.**
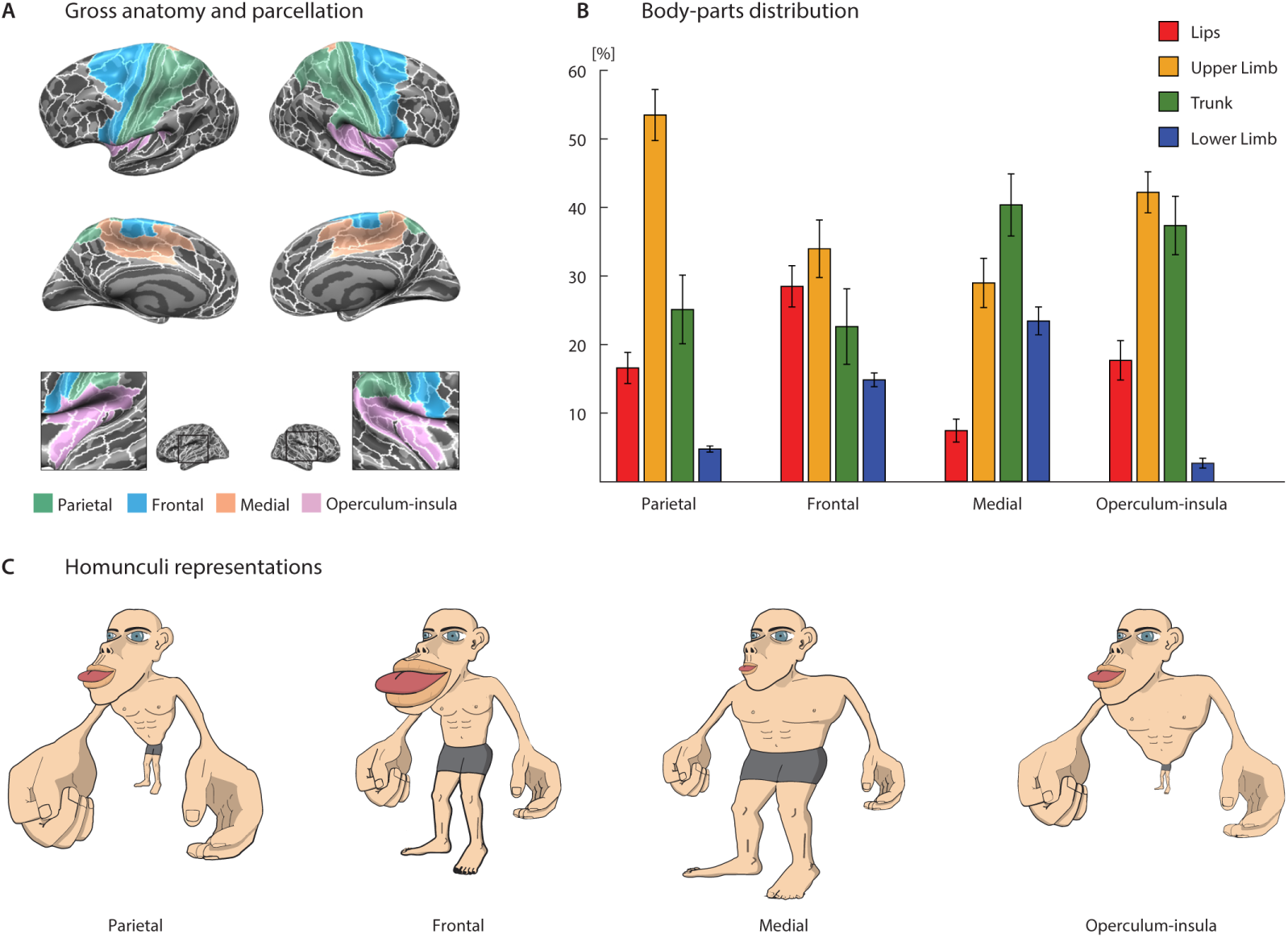
Quantitative differences in the spatial distributions of body-parts (“homunculi”). **(A)** Subdivisions of the somatosensory responsive cortex into four gross anatomical regions: parietal (S1 and posteriorly), frontal (anterior to S1), medial (medial to S1) and operculum-insula (inferior to S1). **(B)** Distribution of the different body-parts [lips, upper limb (proximal and distal combined), trunk (upper and lower combined) and lower limb (proximal, mid and distal combined)] within each gross anatomical region averaged across two hemispheres (Error bars are standard deviation, computed by bootstrapping over participants). **(C)** Homunculi proportional representation of body-parts in each gross anatomical region as modified versions of Penfield’s original homunculus (S1), as in Fig. 1C.

The anterior parietal region, the somatosensory part of the parietal cortex (including S1), was found to encompass 32% of the somatosensory responsive cortex. Somatosensory representation in this region was dominated by the upper limb (53.5±3.8%), more than any other region (p<0.001), including S1. This was followed by a substantial representation of the trunk (25.1±5.1%), which is almost 60% greater than its representation in S1 (p<0.001). The representation of these two body-parts come at the expense of the lips (16.6±2.3%) and the lower limb (4.8±0.5%) (Fig. 2B, see also Fig. S4). In fact, the latter two representations are almost absent outside S1 in the parietal cortex (Fig. 1A and Table S2).

The posterior part of the frontal cortex comprises 18% of the somatosensory responsive cortex. The rank order of body-parts representation in the frontal region is identical to that in S1 (upper limb>lips>trunk>lower limb). However, there are quantitative differences in their relative sizes. The representation of the upper limb (34.0±4.3%) is smaller than its representation in S1 (p<0.001), allowing for a larger representation of the trunk (22.6±5.6%) and lower limb (14.9±1.1%). The relative representation of the lips (28.5 ±3.1%) is comparable to that in S1 (Figs. 2B and S4).

The medial region (12% of the somatosensory responsive cortex) is the most different from S1. The spatial distribution of body-parts in this region is most similar to their veridical proportion (skin area; (Catani, 2017)). The medial region is dominated by the trunk (40.3±4.6%), followed by the upper limb (28.9±3.6%), lower limb (23.4±2.1%) and lips (7.4±1.7%). The trunk and lower limb are more highly represented in the medial region than other regions (p<0.001), whereas the upper limb and lips are the least represented (p<0.001; Figs. 2B and S4).

Finally, the operculum-insular cortex (16% of the somatosensory cortex) is dominated by the upper limb (42.2±3.0%) and trunk (37.4±4.3%). There is less representation of the lips (17.7±2.9%) and the representation of the lower limb is nearly absent (2.7±0.8%; Figs. 2B and S4). We also found somatosensory representations in the area of the temporo-occipital parietal junction and at the inferior-temporal gyrus (22% of the somatosensory responsive cortex). Because these areas are topologically disconnected from S1 along the somatosensory cortex, their analysis is outside the scope of this paper. Taken together, the spatial distributions of body-parts in the four anatomical regions are substantially different from one another, as well as from S1, as represented by the different homunculi corresponding to the four regions (Fig. 2C). These differences may be related to different functional specializations of these regions as discussed below.

## Discussion

Data-driven quantification of the somatosensory system revealed that it comprises a large fraction of the human cortex, which includes high-order regions, suggests that the somatosensory system plays a role in high cognitive processing (which may be entitled “somatosensory cognition”). Quantification of the four gross anatomical regions (parietal, frontal, medial and operculum-insula) revealed that the somatosensory responsive areas differ from each other, and from the primary somatosensory cortex (BAs 1, 2, 3a, 3b). Specifically, they are characterized by different distributions of body-parts representations, manifested in different homunculi (Fig. 2C) that are distinctive from Penfield’s classical S1 homunculus (Fig. 1C). This heterogeneity in the distribution of body-parts representations implies different functional roles of the four homunculi.

The parietal somatosensory region is comprised of S1 and the anterior part of the inferior and superior parietal lobules. Upper limb and trunk representations dominate the parietal somatosensory region. This is particularly pronounced in the anterior parietal lobules (S1 excluded). There, lower limb and lips representation are minimal (table S2). The higher proportion of upper limb and trunk representations in the parietal lobules (compared with S1) is consistent with their role in body-processing related functions such as multisensory integration and bodily self-consciousness (Tsakiris et al., 2007; Kammers et al., 2009; Petkova et al., 2011; Blanke, 2012).

The frontal region includes all vertices that responded to somatosensory stimulation anterior to the central sulcus, including M1 (BA 4), the premotor cortex (BA 6) and parts of the frontal eye field (BA 8). Body-parts representation in M1 is dominated by the lower limb and lips with little representation of upper limb and trunk (table S2). By contrast, lower limb representation is almost absent anterior to M1. This increased upper limb and trunk representation is compatible with high-order functions of this region, since the premotor cortex is important not only for motor planning but also for body agency and embodiment (Blanke, 2012). Electrophysiological studies in non-human primates showed that neurons in the anterior superior parietal lobule and the premotor cortex integrate visual and somatosensory stimuli involving the arm and the trunk (Fogassi et al., 1996; Graziano et al., 1997, 2000; Maravita and Iriki, 2004). Premotor-parietal orchestrated activity was highlighted by functional neuroimaging studies in humans as well as lesion studies in patients demonstrating the putative role of these regions in body processing of the hand and trunk (Blanke, 2012). These results are compatible with the substantial upper limb and trunk representations in both the anterior parietal lobules and the regions anterior to M1 found in our study.

Body-parts distribution in the medial region is closest to human veridical proportions (unlike S1’s “grotesque creature”), as evident in the homunculi illustrations (Fig. 2C). The large representation of the lower limb, the highest of all regions, is consistent with clinical observations of leg-related pathologies (Schneider and Gautier, 1994). Lower limb representation is particularly pronounced in the SMA. As in M1, the dominance of lower limb representation is consistent with the substantial role of the lower limb in motor actions.

Our finding of somatosensory representations in the middle cingulate cortex (MCC) is consistent with previous findings of somatosensory response elicited by electrical stimulation of peripheral nerves (Arienzo et al., 2006) and evoked somatosensory sensation by direct intracranial stimulation of the MCC (Lim et al., 1994). The MCC is important for the conscious perception of pain, as well as the emotional and motivational aspects of pain processing. In particular, the anterior MCC (posterior part of the anterior cingulate cortex, ACC) mediates between emotion and cognition (Vogt, 2005). Finally, this region was also implicated in somatoform disorder (Saj et al., 2014), corroborating its seminal role in body consciousness.

Somatosensory representation was found in the posterior insula which is known to be involved in the processing of body ownership, visceral effects, and emotional modulations (Craig, 2002; Tsakiris et al., 2007; Taylor et al., 2009; Cauda et al., 2012; Grecucci et al., 2013). We find that the representation of the trunk occupies one third of the posterior insula (Table S2). The large proportion of the trunk in this region hints at the trunk’s special role in the emotional aspects of somatosensory processing.

We have also identified somatosensory response in the temporal operculum which includes parts of the auditory cortex. This finding is in agreement with the large body of research regarding multisensory integration (*e.g.*, (Kayser et al., 2005)). Interestingly, somatosensory representation in the associative auditory areas was restricted to the right hemisphere. This asymmetry may be related to the more dominant role of the right hemisphere in auditory spatial processing (Tiitinen et al., 2006).

This study confronts several limitations. First, due to the difficulty in the accessibility within the MRI apparatus and the continuous stimulation applied, we did not cover the entire skin surface of the body. While the classical S1 homunculus presents prominent lips representation, we cannot exclude more prominent representation of upper face/head in homunculi outside of S1, as we have shown for the trunk. Nevertheless, the fact that body-parts spatial distribution in S1 was found comparable to Penfield’s somatosensory homunculus suggests that this did not significantly affected the final results. Further research with a wider MRI machines and additional body coverage may elaborate our findings.

Second, the experimental paradigm investigates whole-brain response to stimulation of each body-part separately, in a continuous manner, and not multiple body-parts or body-sides simultaneously. Further studies may address questions regarding the linearity of the somatosensory response and bilateral interactions as was suggested previously (e.g., Reed et al., 2011).

In conclusion, using a novel approach, the current work has demonstrated how large fraction of the human cortex, including high-order regions, is involved in processing light touch somatosensory stimulation. The spatial distributions of body-parts in different anatomical regions were found to differ substantially from S1. Spatial distributions were unique to each region, and are likely related to the region’s functional specialization. Our findings suggest that somatosensation plays a major role in human cognition. Future research should further develop these findings and incorporate them into a joint framework of “somatosensory cognition”

## General

We thank N. Munro for her important comments on the manuscript and N. Klein for critical assistant with figure graphics. We thank the ELSC MRI unit, Assaf Yohalashet, Lee Ashkenazi and Yuval Porat for their dedicated work.

