## Supporting Information for "The “creatures” of the human cortical somatosensory system"

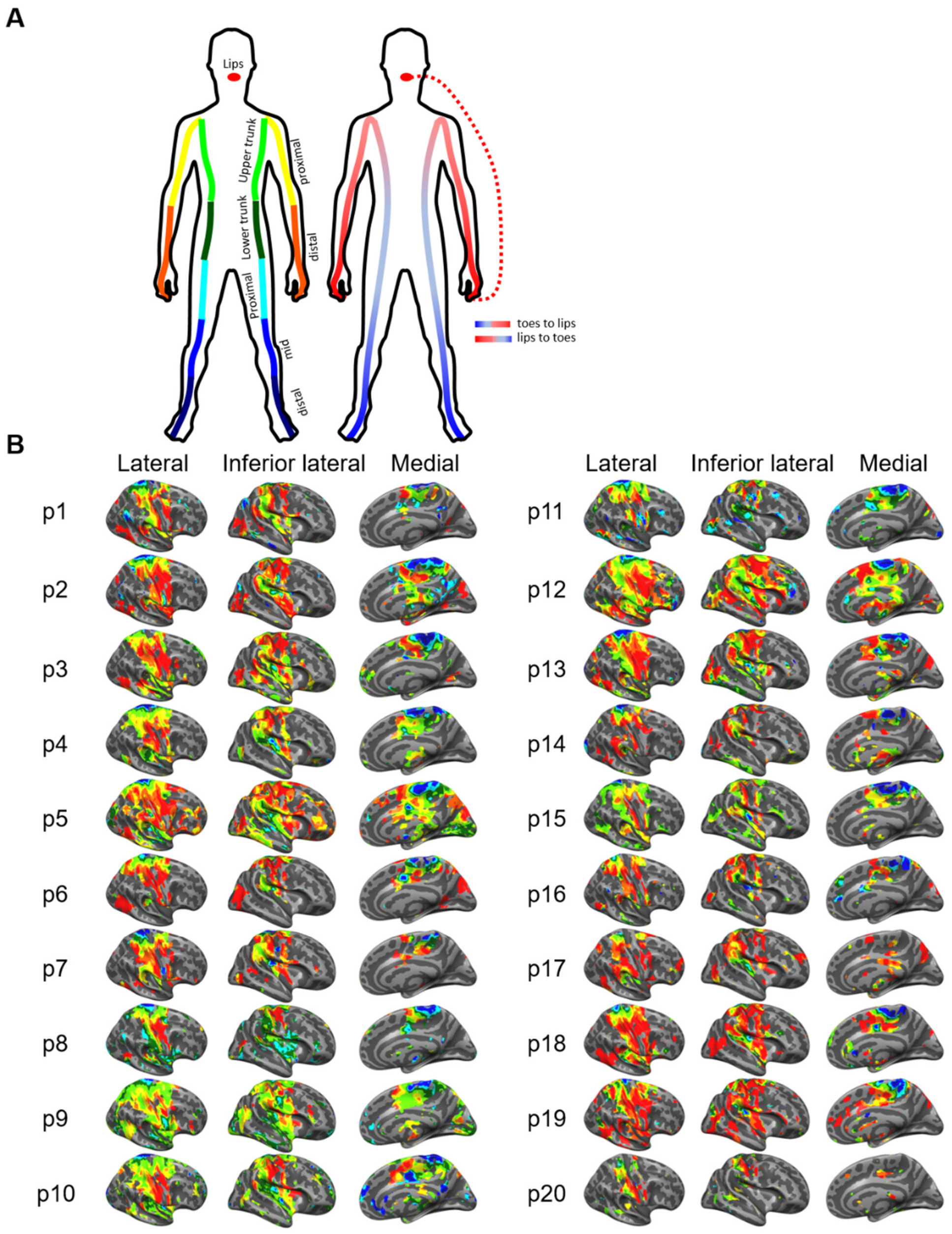


**Fig. S1. Experimental paradigm and cortical distribution of the somatosensory representation-single participants.** (**A**) Scheme of whole-body continuous periodic brush movement from lips-to-toes and from toes-to-lips bilaterally in two different scanning runs (right). Body-parts defined along the continuous stimulation: lips, distal upper limb, proximal upper limb, upper trunk, lower trunk, proximal lower limb, mid lower limb and distal lower limb (left). (**B**) Cross correlation maps corresponding to stimulation of the contralateral body side are shown for single participants (p1-p20) on the lateral, inferior-lateral and medial surfaces of the right hemisphere (t-test, $\mathbf{\alpha = 0.05}$, Bonferroni corrected for multiple correlations; 8 lag values x two paradigms, 134 degrees of freedom, $p<0.003$). Preferred body part is the lag with the maximum value of averaged correlation distributions of both start lips and start toes directions. Color code represents body-parts as depicted in the scheme above.

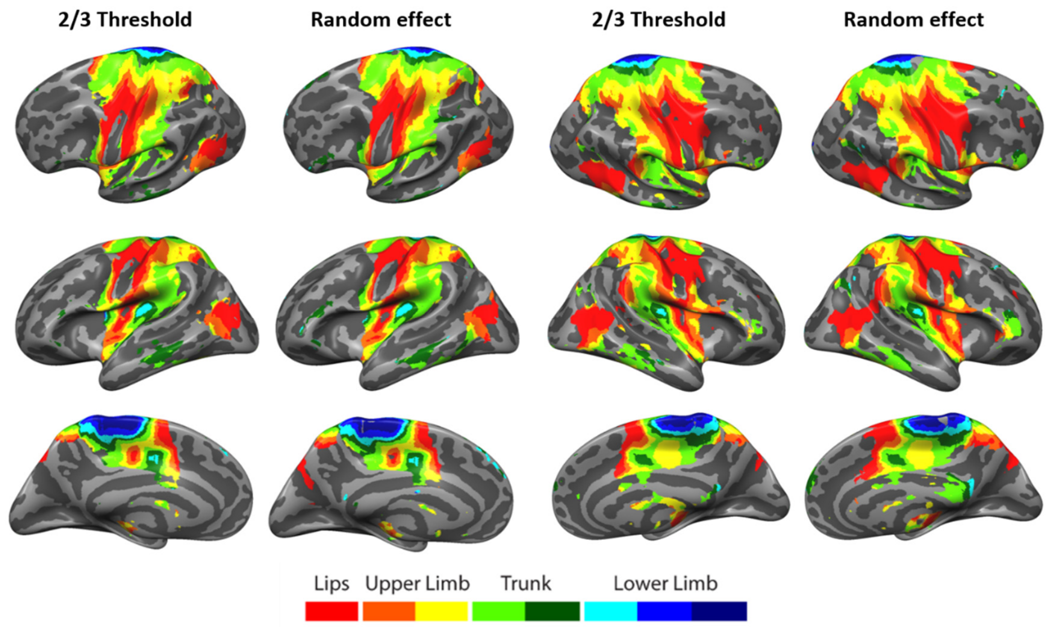


**Fig. S2. Somatosensory cortical representation- comparison between set-threshold and random effect for group significance.** Cross correlation group maps (n=20) corresponding to stimulation of the contralateral body side are shown on the lateral, inferior-lateral and medial surfaces. Significance was computed in two methods: set-threshold (2/3 Threshold) and random effect (RE). Random effect computation: correlation coefficients in each vertex were averaged between two stimuli directions (lips to toes and the opposite). The 8 resulting correlation coefficients (corresponding to cross correlations with 8 lags) were transformed to t values by: $\mathbf{t = r \cdot \sqrt{\frac{n-2}{1-r^2}}}$ where n is the sample size (number of measurements-TR). We then applied a one tailed t-test ($\mathbf{H_0: \mu = 0, H_1: \mu > 0}$) on the t-values from all participants for each lag separately. Significance threshold was set to $\mathbf{\alpha = 0.05}$ to the smallest p value from all lags, Bonferroni-corrected for 8 comparisons. These spatial distribution maps are qualitatively similar using the two methods.

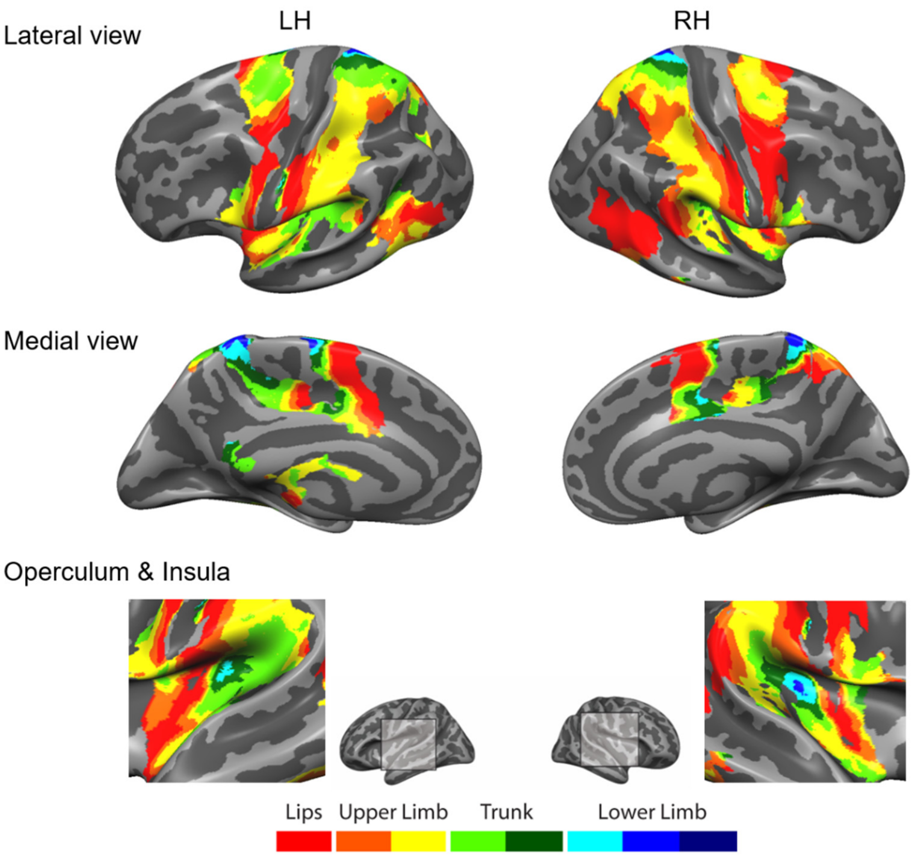


**Fig. S3. Ipsilateral cortical distribution of the somatosensory representation.** Cross correlation maps corresponding to stimulation of the ipsilateral body side are shown on the lateral and medial surfaces as well as the opercular and insular cortex (LH: left hemisphere; RH: right hemisphere; level significance- at least 2/3 of the participants with significance $p<0.05$). Color code represents body-parts: lips (red), distal upper limb (orange), proximal upper limb (yellow), upper trunk (light green), lower trunk (green), proximal lower limb (light blue), mid lower limb (blue) and distal limb (dark blue).

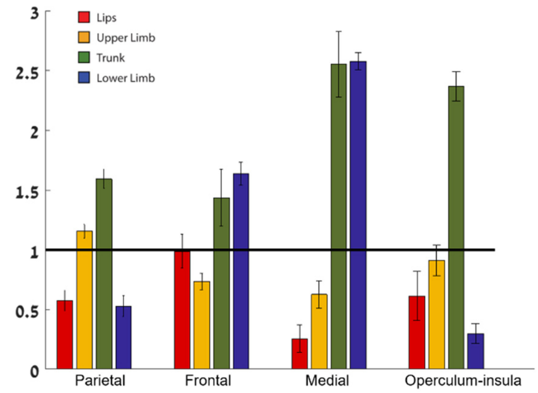


**Fig. S4. Cortical representation of body parts relative to S1.** The ratio between the number of vertices within each of the gross anatomical regions responding to each body part [lips, upper limb (proximal and distal combined), trunk (upper and lower combined) and lower limb (proximal, mid and distal combined)] and the number of vertices of the same body part within S1 (BAs 3a,3b,1 and 2).

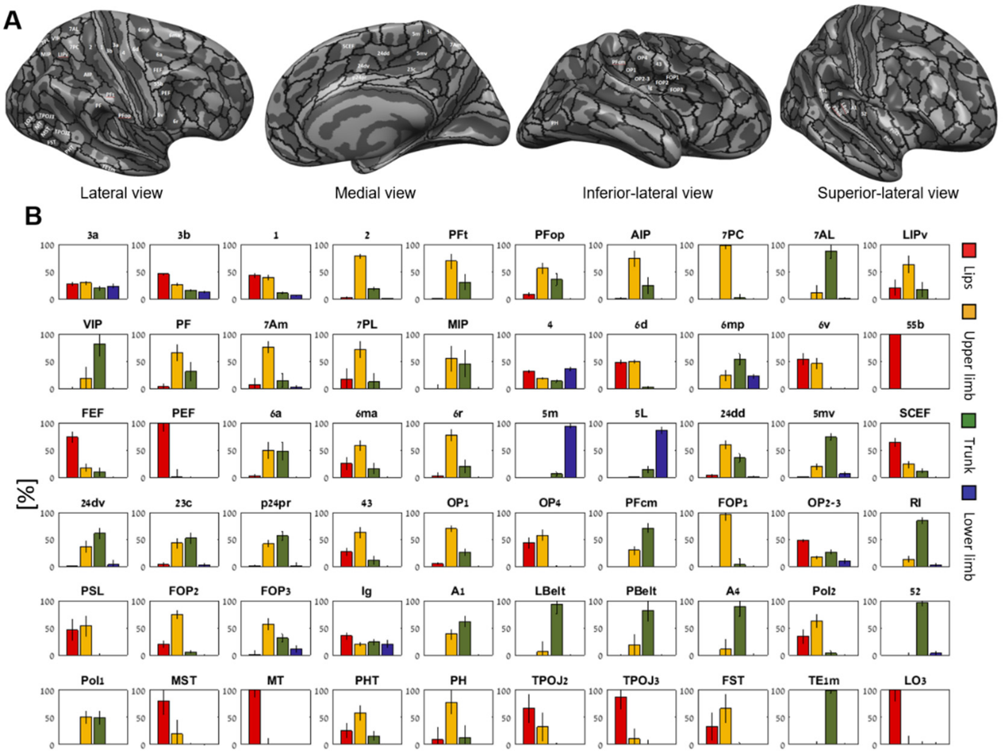


**Fig. S5. Somatosensory responsive parcellation areas- location and body-parts spatial distribution.** (**A**) Multi-modal data-driven cortical parcellation by Glasser et al. (2016) delineate 180 areas per hemisphere presented on an averaged inflated cortical surface of the right hemisphere (FreeSurfer's, fsaverage template brain; (Desikan et al., 2006)) in four different views. 60 parcellation areas were found responsive to somatosensory stimulation (named in white), 50 of them were found in the left and right hemispheres, 9 unique to the right hemisphere and 1 to the left hemisphere (LO3). (**B**) Distribution of the different body-parts [lips, upper limb (proximal and distal combined), trunk (upper and lower combined) and lower limb (proximal, mid and distal combined)] within each somatosensory responsive parcellation area. Percentages of body parts within homologues areas in both right and left hemispheres were averaged (errobars, bootstrapping over participants with 1,000 iterations).

**Table S1. Somatosensory responsive parcellation areas.** 50 Somatosensory responsive parcellation areas were found in the left (L) and right (R) hemispheres, additional 9 unique to the right hemisphere and 1 to the left hemisphere. Somatosensory responsive areas are defined as areas that contain more than 50% of vertices responding significantly (cross correlation, p<0.05, group level) to contralateral body stimulation. The number of significant vertices out of the total number within an area is presented. Areas were classified into four gross anatomical regions (R-region; P-parietal, F-frontal, M-medial and OI-operculum-insula). Body-parts spatial distribution for all areas and both hemispheres is presented by the percentage of the lips, hand, trunk and leg (errors in brackets, bootstrapping over participants with 1,000 iterations).

| Area | H | Significant vertices (R\|L) | | R | Lips (R\|L) [%] | | Upper limb (R\|L) [%] | | Trunk (R\|L) [%] | | Lower limb (R\|L) [%] | |
|---|---|---|---|---|---|---|---|---|---|---|---|---|
| **3a** | R,L | 1401/1770 | 1186/1812 | P | 30(6) | 24(5) | 25(5) | 33(5) | 19(4) | 20(6) | 25(5) | 22(4) |
| **3b** | R,L | 2518/2552 | 2680/2845 | P | 47(3) | 44(3) | 23(2) | 30(3) | 16(2) | 14(2) | 13(1) | 12(1) |
| **1** | R,L | 1998/2045 | 2124/2162 | P | 45(7) | 43(5) | 38(7) | 41(5) | 12(2) | 9(1) | 5(1) | 7(1) |
| **2** | R,L | 2913/2913 | 2554/2554 | P | 2(4) | 3(3) | 81(5) | 76(4) | 17(2) | 20(3) | 1(0) | 1(1) |
| **PFt** | R,L | 1083/1083 | 1205/1260 | P | 0(3) | 0(1) | 90(11) | 49(22) | 10(12) | 51(22) | 0(0) | 0(0) |
| **PFop** | R,L | 735/735 | 839/839 | P | 10(6) | 6(3) | 60(9) | 52(16) | 29(10) | 42(16) | 0(0) | 0(0) |
| **AIP** | R,L | 1472/1710 | 1044/1639 | P | 2(4) | 0(4) | 79(17) | 70(21) | 19(17) | 30(22) | 0(0) | 0(0) |
| **7PC** | R,L | 1010/1010 | 865/865 | P | 0(3) | 0(3) | 98(5) | 98(13) | 2(4) | 2(13) | 0(0) | 0(0) |
| **7AL** | R,L | 910/910 | 1082/1082 | P | 0(0) | 0(1) | 16(14) | 7(17) | 82(14) | 92(16) | 2(2) | 1(2) |
| **LIPv** | R,L | 469/661 | 578/765 | P | 13(25) | 27(20) | 55(22) | 71(19) | 32(23) | 2(17) | 0(0) | 0(0) |
| **VIP** | R,L | 525/525 | 699/699 | P | 0(4) | 0(3) | 11(26) | 26(27) | 89(27) | 74(28) | 0(0) | 0(0) |
| **7Am** | R,L | 973/1061 | 781/1047 | P | 4(10) | 10(18) | 79(13) | 73(16) | 15(16) | 14(15) | 1(1) | 4(6) |
| **PF** | R | 1454/1681 | - | P | 3(6) | - | 66(16) | - | 31(17) | - | 0(0) | - |
| **7PL** | R,L | 354/429 | 426/531 | P | 5(15) | 29(31) | 82(16) | 60(24) | 13(18) | 11(19) | 0(0) | 0(0) |
| **MIP** | R | 618/815 | - | P | 0(8) | - | 55(23) | - | 45(24) | - | 0(2) | - |
| **4** | R,L | 3187/3855 | 3127/4015 | F | 34(4) | 29(4) | 16(2) | 21(2) | 16(3) | 11(3) | 34(4) | 39(4) |
| **6d** | R,L | 876/876 | 917/917 | F | 55(6) | 41(5) | 45(6) | 55(6) | 0(0) | 4(4) | 0(0) | 0(0) |
| **6mp** | R,L | 1454/1454 | 1434/1434 | F | 0(0) | 0(0) | 35(13) | 13(12) | 53(13) | 55(12) | 12(3) | 32(5) |
| **6v** | R,L | 694/709 | 425/463 | F | 56(10) | 52(16) | 44(9) | 48(14) | 0(3) | 0(5) | 0(0) | 0(0) |
| **FEF** | R,L | 830/850 | 672/832 | F | 96(8) | 52(18) | 4(7) | 30(10) | 0(0) | 19(14) | 0(0) | 0(0) |
| **6a** | R,L | 1334/1715 | 1441/1765 | F | 5(6) | 0(3) | 59(13) | 40(22) | 36(16) | 60(23) | 0(0) | 0(0) |
| **6ma** | R,L | 739/1217 | 497/966 | F | 35(13) | 15(15) | 48(9) | 69(14) | 16(12) | 16(11) | 0(0) | 0(1) |
| **6r** | R,L | 683/1287 | 855/1176 | F | 5(10) | 0(6) | 93(12) | 63(18) | 2(12) | 37(19) | 0(0) | 0(0) |
| **55b** | R | 315/421 | - | F | 100(1) | - | 0(1) | - | 0(0) | - | 0(0) | - |
| **PEF** | R | 404/480 | - | F | 100(16) | - | 0(16) | - | 0(1) | - | 0(0) | - |
| **5m** | R,L | 789/805 | 503/503 | M | 0(1) | 1(1) | 19(6) | 21(8) | 75(7) | 73(8) | 6(4) | 5(5) |
| **5L** | R,L | 904/904 | 854/854 | M | 0(0) | 0(0) | 0(2) | 0(1) | 11(7) | 17(9) | 89(7) | 83(9) |
| **24dd** | R,L | 990/990 | 1115/1115 | M | 3(3) | 4(4) | 59(7) | 60(0) | 37(8) | 34(11) | 0(1) | 2(2) |
| **5mv** | R,L | 1162/1243 | 932/977 | M | 0(1) | 1(1) | 19(6) | 21(8) | 75(7) | 73(8) | 6(4) | 5(5) |
| **SCEF** | R,L | 525/875 | 674/1139 | M | 69(9) | 60(11) | 17(7) | 31(9) | 14(6) | 9(4) | 0(1) | 0(1) |

| | | | | | | | | | | | | |
|---|---|---|---|---|---|---|---|---|---|---|---|---|
| **24dv** | R,L | 752/752 | 677/677 | M | 0(2) | 0(3) | 39(14) | 34(12) | 61(14) | 60(12) | 0(7) | 6(13) |
| **23c** | R,L | 792/1325 | 979/1253 | M | 0(2) | 6(7) | 48(14) | 38(11) | 52(14) | 51(11) | 0(1) | 5(4) |
| **p24pr** | R,L | 556/556 | 450/452 | M | 2(2) | 0(3) | 47(8) | 36(8) | 50(9) | 62(10) | 0(4) | 2(10) |
| **43** | R,L | 713/748 | 527/563 | OI | 50(15) | 4(7) | 50(14) | 75(14) | 1(4) | 21(15) | 0(0) | 0(0) |
| **OP1** | R,L | 468/468 | 760/760 | OI | 0(3) | 0(1) | 100(6) | 94(21) | 0(5) | 6(21) | 0(0) | 0(1) |
| **OP4** | R,L | 1045/1096 | 898/902 | OI | 33(11) | 53(13) | 67(11) | 47(13) | 0(1) | 0(2) | 0(0) | 0(0) |
| **PFcm** | R,L | 708/708 | 605/679 | OI | 0(1) | 0(0) | 35(9) | 24(11) | 65(9) | 76(11) | 0(0)) | 0(0) |
| **FOP1** | R,L | 249/325 | 185/299 | OI | 0(3) | 0(1) | 100(6) | 94(21) | 0(5) | 6(21) | 0(0) | 0(1) |
| **OP2-3** | R,L | 568/568 | 665/665 | OI | 43(3) | 52(4) | 20(3) | 13(3) | 31(4) | 22(3) | 5(5) | 13(4) |
| **RI** | R,L | 847/847 | 859/909 | OI | 0(0) | 0(1) | 21(7) | 5(7) | 77(7) | 91(9) | 2(2) | 4(5) |
| **FOP2** | R,L | 411/436 | 400/400 | OI | 30(12) | 9(7) | 70(12) | 79(8) | 0(0) | 12(6) | 0(0) | 0(0) |
| **FOP3** | R,L | 243/450 | 328/516 | OI | 1(18) | 0(1) | 99(18) | 15(11) | 0(9) | 63(11) | 0(1) | 23(12) |
| **Ig** | R,L | 436/489 | 449/544 | OI | 38(8) | 33(10) | 22(5) | 19(5) | 28(6) | 20(6) | 12(6) | 28(8) |
| **A1** | R,L | 295/301 | 395/400 | OI | 0(1) | 0(0) | 43(13) | 34(10) | 57(13) | 66(10) | 0(0) | 0(1) |
| **PSL** | R | 714/877 | - | OI | 46(20) | - | 54(19) | - | 0(3) | - | 0(0) | - |
| **LBelt** | R | 386/457 | - | OI | 0(0) | - | 6(19) | - | 94(19) | - | 0(0) | - |
| **PBelt** | R | 411/499 | - | OI | 0(2) | - | 19(19) | - | 81(19) | - | 0(1) | - |
| **A4** | R | 509/855 | - | OI | 0(1) | - | 10(19) | - | 90(18) | - | 0(2) | - |
| **Pol1** | R,L | 844/867 | 732/774 | OI | 0(0) | 0(3) | 39(15) | 62(12) | 61(15) | 38(13) | 0(1) | 0(1) |
| **Pol2** | R,L | 749/817 | 643/805 | OI | 38(19) | 31(14) | 57(16) | 68(14) | 5(5) | 2(6) | 0(0) | 0(0) |
| **52** | R,L | 316/386 | 319/507 | OI | 0(0) | 0(0) | 0(8) | 0(2) | 95(9) | 98(5) | 5(2) | 2(4) |
| **MST** | R,L | 330/385 | 351/360 | - | 100(31) | 62(31) | 0(30) | 38(30) | 0(2) | 0(5) | 0(0) | 0(0) |
| **MT** | R,L | 201/303 | 268/292 | - | 100(14) | 100(16) | 0(14) | 0(15) | 0(1) | 0(2) | 0(0) | 0(0) |
| **TPOJ2** | R,L | 821/827 | 396/597 | - | 99(30) | 36(33) | 1(30) | 64(31) | 0(0) | 0(5) | 0(0) | 0(0) |
| **TPOJ3** | R,L | 369/532 | 365/400 | - | 81(30) | 96(26) | 17(22) | 4(26) | 2(12) | 0(2) | 0(1) | 0(0) |
| **TE1m** | R,L | 552/639 | 524/663 | - | 0(0) | 0(0) | 0(10) | 0(2) | 100(10) | 100(4) | 0(1) | 0(3) |
| **FST** | R,L | 317/401 | 459/550 | - | 56(36) | 12(26) | 44(36) | 88(26) | 0(3) | 0(6) | 0(0) | 0(0) |
| **PHT** | R | 635/759 | - | - | 26(14) | - | 58(13) | - | 16(8) | - | 0(1) | - |
| **PH** | R | 386/720 | - | - | 9(23) | - | 77(21) | - | 13(22) | - | 0(0) | - |
| **LO3** | L | - | 251/325 | - | - | 100(20) | - | 0(16) | - | 0(6) | - | 0(3) |

**Table S2. Spatial distribution of body-parts.** Percentage of body-parts (lips, upper limb, trunk and lower limb) by anatomical area. Errors were computed by bootstrapping over participants with 1,000 iterations.

| | **Lips** | **Upper Limb** | **Trunk** | **Lower limb** |
|---|---|---|---|---|
| **Somatosensory cortex** | 20.1±3.3 | 41.1±3.3 | 31.1±5.3 | 7.6±0.9 |
| **Parietal** | 16.6±2.3 | 53.5±3.8 | 25.1±5.1 | 4.8±0.5 |
| **Frontal** | 28.5±3.1 | 34.0±4.3 | 22.6±5.6 | 14.9±1.1 |
| **Medial** | 7.4±1.7 | 28.9±3.6 | 40.3±4.6 | 23.4±2.1 |
| **Operculum-insula** | 17.7±2.9 | 42.2±3.0 | 37.4±4.3 | 2.7±0.8 |
| **S1** | 28.9±2.6 | 46.3±2.5 | 15.8±1.3 | 9.1±0.8 |
| **M1** | 31.4±3.5 | 18.7±1.6 | 13.7±2.3 | 36.2±3.2 |
| **Parietal,S1 excluded** | 4.1±3.1 | 60.5±8.2 | 34.9±9.8 | 0.4±0.4 |
| **Frontal, M1 excluded** | 27.0±3.7 | 41.0±5.9 | 27.0±7.7 | 4.9±0.7 |
| **Insular cortex** | 16.3±4.4 | 46.3±4.0 | 32.6±4.9 | 4.8±1.3 |
